# A tail of plumage colouration: disentangling geographic, seasonal, and dietary effects on plumage colour in a migratory songbird

**DOI:** 10.1101/2021.07.09.451669

**Authors:** Sean M. Mahoney, Matthew W. Reudink, Andrea Contina, Kelly A. Roberts, Veronica T. Schabert, Emily G. Gunther, Kristen M. Covino

## Abstract

Plumage ornamentation in birds serves critical inter- and intra-sexual signaling functions. While carotenoid-based plumage colouration is often viewed as a classic condition-dependent sexually selected trait, plumage colouration can be influenced by a wide array of both intrinsic and extrinsic factors. Understanding the mechanisms underlying variation in colouration is especially important for species where the signaling function of ornamental traits is complex or the literature conflicting. Here, we examined variation in the yellow/orange tail feathers of American redstarts (*Setophaga ruticilla*) passing through two migratory stopover sites in eastern North America during both spring and fall migration to assess the role of geographic variation and seasonality in influencing feather colouration. In addition, we investigated whether diet during moult (inferred via stable isotope analysis of feather δ^15^N and δ^13^C) influenced plumage colouration. Our findings indicate that geographic variation, season, and diet all influence individual differences in American redstart colouration, represented by both traditional and tetrahedral colour variables. The extent to which these factors influence colour expression however is largely dependent on the colour metric under study, likely because different colour metrics reflect different attributes of the feather (e.g., structural components vs. pigment deposition). The effects of diet (δ^15^N) and season were pronounced for brightness, suggesting a strong effect of diet and feather wear/degradation on feather structure. Though hue, a metric that should strongly reflect pigment deposition, also changed from spring to fall, that effect was dependent on age, with only adults experiencing a reduction in ornamentation. Taken together, our results highlight the numerous sources of variation behind plumage coloration and underscores the difficulty of unraveling complex visual signaling systems, such as those in American redstarts.

## Introduction

The colourful plumage exhibited by birds serves important roles for both inter-and intrasexual communication (reviewed in Hill 2006; Santos et al. 2011). The literature linking variation in plumage colouration to individual quality and reproduction is multitudinous (Lope and Møller 1993; Griffith and Pryke 2006; Hill 2006; Henschen et al. 2017; Ferree 2019; LaBarbera et al. 2019; Enbody et al. 2018), yet the patterns are often complex and may vary across seasons and populations (Marini et al. 2015; Reudink et al. 2015). As such, much research has been dedicated to understanding the mechanisms underlying variation in the expression of ornamental plumage colouration (Siefferman and Hill 2005; Griggio et al. 2009; Newton and Dawson 2011; Doutrelant et al. 2012; Reudink et al. 2015).

Plumage colouration in birds can be produced by various mechanisms, including the deposition of pigments (e.g., carotenoids, melanins, and porphyrines) and the arrangement of melanosomes within the feather microstructure producing structural (purple, blue, green, and iridescent) colours. Carotenoid colouration (yellow, orange, red plumage) has received by far the most attention, in large part because the expression of carotenoid-based colouration is directly dependent on birds acquiring carotenoids in the diet (the pigments cannot be synthesized *de novo*) and subsequently modifying dietary pigments through a costly physiological process (Lozano et al. 1994; Hill and Montgomerie 1994; Weaver et al. 2018). Thus, carotenoid-based colouration is often viewed as a classic honest indicator of individual condition and quality (Hill 2011; Hill and Johnson 2012; Weaver et al. 2018; but see Simons et al. 2015).

Diet during feather growth has been directly linked to ornamentation and feather quality in both experimental (Hill 2000; Peters et al. 2008; Koch et al. 2017) and observational studies (Slagsvold and Lifjeld 1985; Arriero and Fargallo 2006; Eeva et al. 2009; but see Ferns and Hinsley 2008). For observational studies of free-living birds, one of the challenges is determining dietary conditions during moult (Besozzi et al. 2021). However, stable isotopes of carbon (δ^13^C) and nitrogen (δ^15^N) vary predictably in the environment and are incorporated into growing tissues (Fry 2006; Bowen 2010). Because feathers are inert once grown, they can provide insight into diet during tissue growth. δ^13^C has been used extensively in dietary studies because carbon isotope ratios vary based on plant water stress and photosynthetic system (Lajtha and Marshall 1994). Further, since these isotope elements are transferred up the food chain and eventually incorporated into animal tissues, they can provide important insight into habitat and diet during tissue growth (Hobson 1999). Nitrogen is commonly employed in food web studies, as ^15^N is preferentially incorporated into animal tissues as one moves up the food chain (Post 2002; Poupin et al. 2011). Thus, analysis of δ^15^N tissue values can provide insight into the trophic level of animals within the environment. However, δ^15^N values can also be influenced by environmental conditions, as they tend to be positively associated with temperature and negatively associated with rainfall (Craine et al. 2009).

A recent study on Bullock’s orioles (*Icterus bullockii*) demonstrated that feather δ^15^N values were directly associated with feather carotenoid content and hue (Sparrow et al. 2017). In the case of Bullock’s orioles, birds with lower feather δ^15^N values expressed more orange-shifted hue. Because Bullock’s orioles rely on both insects and fruit during moult, the authors suggested that individuals with a relatively higher amount of fruit in their diet (or those moulting in habitats that resulted in lower δ^15^N values) may have been in better condition and better able to metabolically convert dietary carotenoids to red or orange keto-carotenoids (McGraw 2006; Sparrow et al. 2017; Weaver et al. 2018).

Though feathers are metabolically inert once grown, feather colours are not necessarily static across the life of the feather. Feather wear can modify feather colouration; for example, European starlings (*Sturnus vulgaris*) lose their distinctive winter spots through wear and abrasion on the feather tips leaving only the melanin-based feather structure behind for the breeding season. For most species, these changes are less dramatic but can still have important fitness consequences. Feather abrasion, ultraviolet (UV) radiation, and feather degrading bacteria can all lead to changes in ornamentation after initial feather growth (Örnberg et al. 2002; Shawkey and Hill 2004). For example, carotenoid-based red colouration in House finches (*Haemorhous mexicanus*) becomes increasingly more saturated and hue becomes more yellow-shifted in the spring following moult (McGraw and Hill 2004). Melanin-based plumage in Fox sparrows (*Passerella iliaca*) also changes colour over time (Weckstein et al. 2002), possibly due to UV exposure or accumulation of dirt and oils on the feathers (Montgomerie 2006). Additionally, colour in many species changes with age — sometimes dramatically through delayed plumage maturation (Hawkins et a. 2012) and sometimes more subtly as individuals age (Marini et al. 2015; Ward et al. 2021); therefore, it is important for studies to assess the environmental, biological, ontological, and temporal influences on feather colour to help unravel the mechanisms regulating this phenotypic trait.

For the American redstart (*Setophaga ruticilla*), the brilliant orange and black plumage ornamentation expressed by males in definitive plumage has been extensively studied (e.g., Reudink et al. 2009a,b; Marini et al. 2015; Reudink et al. 2015), yet the role of plumage for inter- and intra-sexual signaling remains frustratingly complicated and unresolved (see Marini et al. 2015; Table 1). As such, recent research on this species has focused on understanding the mechanisms underlying variation in plumage colouration (Marini et al. 2015; Reudink et al. 2015). In an 11-year study, Reudink et al. (2015) found that rainfall and temperature during moult was directly linked variation in feather hue; however, individuals themselves also appear to change throughout life, with males expressing the most orange-shifted hue during their first breeding season in definitive plumage (Marini et al. 2015). At a broader scale, American redstarts also exhibit marked geographic variation in plumage ornamentation across the range (Norris et al. 2007). Adding to the complexity of this system, American redstarts exhibit delayed plumage maturation, with young (hatch-year [HY] and second-year [SY]) males exhibiting female-like grey and yellow plumage throughout their first year of life, while adult (after hatch year [AHY] and after second year [ASY]) birds display jet black and orange plumage. As such, male redstarts of different age classes appear to utilize different metabolic pathways prior to depositing ingested dietary carotenoids. Adding to the complexity, these potentially important signals are grown at different times of year and under different conditions — in the nest in early-to-mid summer for young birds and after breeding in late summer for adult birds. Thus, American redstarts in different age- and sex-classes are likely differentially affected by both intrinsic and extrinsic factors.

**Table 1.** Linear mixed effects model selection results using a sample-size corrected Akaike’s Information Criterion (AICc) method. Models were built to predict the effects of American redstart colouration (traditional colourimetric values (from Montgomerie 2006): brightness, red chroma, red hue and tetrahedral colourspace values (from Stoddard and Prum 2008): hue θ, hue Φ, r.achieved, and luminance) based on sex, site, season, age, δ^15^N, and δ^13^C. Final models used in analyses are indicated in bold. ΔAIC: Change between full and reduced model; AICcWt: AICc weights, indicating the probability a model is the most parsimonious model; LL: Log-likelihood.

| Variable | Model | K | AICc | $\Delta\text{AICc}$ | AICcWt | LL |
| --- | --- | --- | --- | --- | --- | --- |
| Brightness | **Reduced** | **6** | **-473.7** | **0** | **0.8** | **243.2** |
|  | Full | 9 | -470.8 | 3 | 0.2 | 245.1 |
| Red Chroma | **Reduced** | **4** | **-537.5** | **0** | **1** | **272.9** |
|  | Full | 11 | -529 | 8.5 | 0 | 276.6 |
| Red Hue | **Reduced** | **7** | **79.3** | **0** | **0.9** | **-32.2** |
|  | Full | 11 | 84.4 | 5.1 | 0.1 | -30.1 |
| Hue $\theta$ | **Reduced** | **9** | **-327.1** | **0** | **0.9** | **173.3** |
|  | Full | 11 | -322.5 | 4.7 | 0.1 | 173.3 |
| Hue $\Phi$ | **Reduced** | **7** | **-93.9** | **0** | **0.9** | **54.4** |
|  | Full | 11 | -88.9 | 5.0 | 0.1 | 56.5 |
| r.achieved | **Reduced** | **6** | **-424.2** | **0** | **0.8** | **218.4** |
|  | Full | 11 | -421.6 | 2.6 | 0.2 | 222.9 |
| Luminance | **Reduced** | **4** | **-444.5** | **0.0** | **0.9** | **226.4** |
|  | Full | 11 | -440.3 | 4.2 | 0.1 | 232.2 |

Our goal was to examine the relative importance of the several factors that may underlie variation in the expression of ornamental colouration in the American redstart. We investigated plumage colouration of American redstarts passing through two different migration stopover sites in eastern North America (Fig. 1, Braddock Bay Bird Observatory (BBBO) and Appledore Island Migration Station (AIMS)) during both spring and fall migration and examined the stable isotope ratios of δ^13^C and δ^15^N in the feathers of each individual. As such, we were able to concurrently determine the relative effects of geographic (site), temporal (season), biological (sex), ontological (age), environmental and dietary variation (δ^13^C and δ^15^N), along with age and sex, in the expression of plumage ornamentation. We predicted there would be differences in plumage colouration between sites and that birds captured in fall would exhibit a greater degree of ornamentation (increased red chroma, orange-shifted hue, higher brightness) compared to those captured in spring. In addition, we predicted that birds with more negative δ^13^C feather isotope values (indicating habitats with lower water stress) and more positive δ^15^N feather isotope values (indicating feeding at a higher trophic level) would exhibit increased ornamentation.

**Fig. 1.**
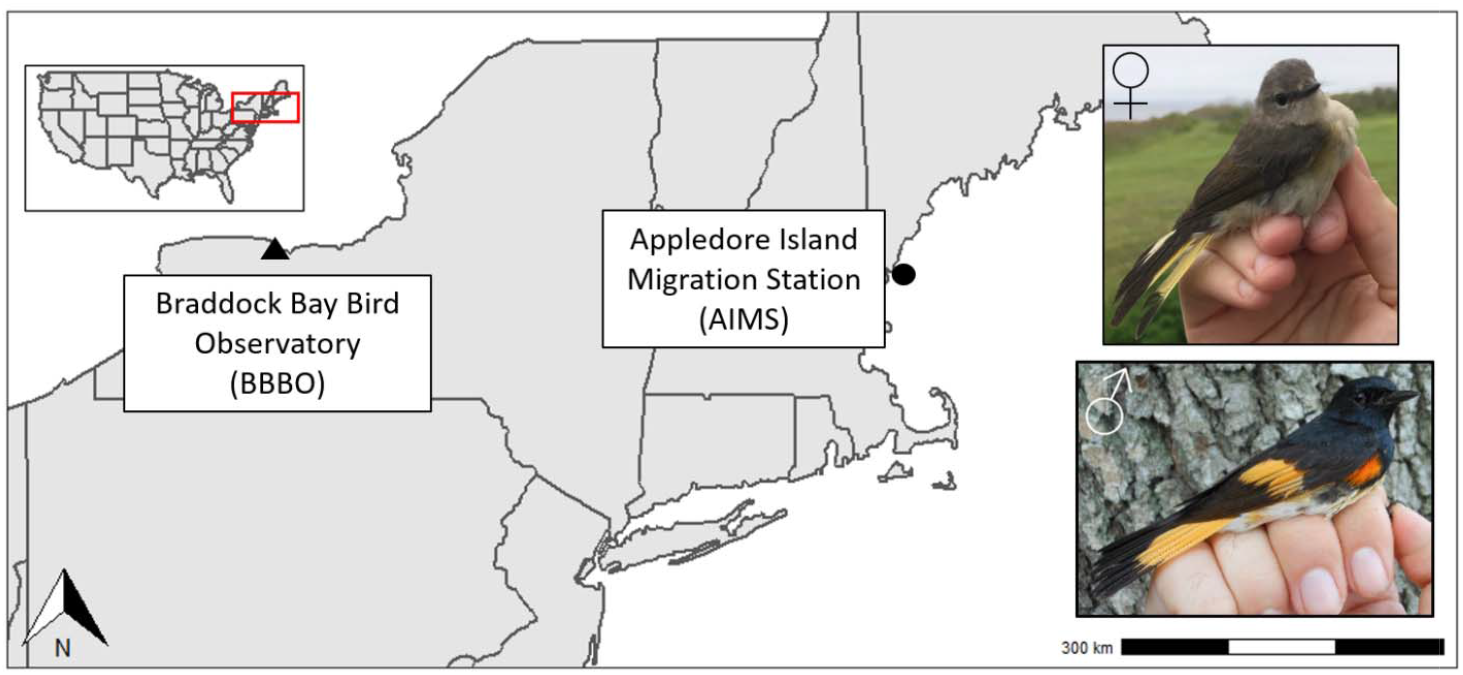
Map showing study sites in New York, US (Braddock Bay Bird Observatory) and off the coast of Maine, US (Appledore Island Migration Station). Inset photos show adult female (top) and adult male (bottom) American redstarts (photo credit: Kristen M. Covino).

## Methods

### Field Methods

We collected feather samples from American redstarts, a sexually dimorphic Nearctic-Neotropical migrant that breeds in deciduous forests in the United States and Canada (Sherry et al. 2020). Our collection sites were two locations of similar latitude but different proximity to the Atlantic coast: Braddock Bay Bird Observatory (43° 9’N, 77° 36’W) in New York State and Appledore Island Migration Station (42°58N, 70°36’ W) off the coast of Maine. Braddock Bay Bird Observatory (BBBO) is an inland site just south of Lake Ontario and the Appledore Island Migration Station (AIMS) is located seven miles offshore from the Maine-New Hampshire border (Fig. 1).

Field researchers captured birds at AIMS using mist nets (weather permitting) checked at least every 20 minutes from sunrise to sunset from 11 May to 7 June during spring migration and 14 August to 15 September during fall migration 2016 and 2017. During 2016 and 2017 at BBBO field researchers captured birds with mist nets checked every half hour starting at sunrise and for 6 hours thereafter from 14 April through 8 June (spring) and 12 August through 2 November (fall). A United States Geological Service leg band was placed on each bird and sex and age were determined according to Pyle (1997) based on plumage, molt, and/or degree of skull pneumatization. In fall, age was determined to be hatch-year (hereafter young) or after-hatch-year (hereafter adult). In spring, age was determined to be second-year (young) or after-second-year (adult). We sampled the left and right fifth rectrix feathers (second from the outside on each side of the tail), provided the feathers were not replaced outside of regular molt, in which case we did not sample feathers and that individual was excluded from this study. In total, we collected feathers from 134 individual American redstarts; 59 from BBBO and 75 from AIMS (Table S1).

### Sample preparation and stable isotope analysis

We conducted analysis of stable isotope ratios of carbon (^13^C/^12^C) and nitrogen (^15^N/^14^N) on keratin tissues obtained from feathers plucked during mist-netting efforts at AIMS and BBBO. We cleaned each feather to remove oils and debris with immersion in a 2:1 chloroform-methanol solution followed by air-drying for 24-h under a fume hood. We then washed each feather with a 1:30 solution of detergent (Fisher Scientific Versa-Clean #04-0342) and deionized water before rinsing three times with deionized water (Paritte and Kelly 2009; Chew et al. 2019). We cut a 350 μg (± 10 μg) fragment and packed in tin capsules placed in a Carlo Erba NC2500 elemental analyzer (CE Instruments, Milano, Italy), interfaced with a Thermo Delta V+ isotope ratio mass spectrometer (Thermo-Finnigan, Bremen, Germany) operated by the Central

Appalachians Stable Isotope Facility (CASIF) at the Appalachian Laboratory (Frostburg, Maryland, USA). We matched each sample with powdered keratin (porcine) from Spectrum Chemicals, a quality control (QC) in-house standard. We report feather carbon (δ^13^C) and nitrogen (δ^15^N) as mean ± SD in delta notation of parts per thousand (‰) from the standards (δD_sample_ = [(*R*_sample_/*R*_standard_) - 1] × 1,000) compared to the Vienna PeeDee Belemnite (V-PDB) and AIR scales, respectively. The standard deviation of the reference material was ±0.12‰ (δ^13^C) and ±0.11‰ (δ^15^N).

### Colour Variation

Prior to cutting and packaging feathers (above) but after the above-described cleaning steps, feather colouration was determined by measuring the reflectance within the avian visual range (300-700nm) with an Ocean Optics Jaz Spectrometer (Reudink et al. 2015). The yellow-orange vane sections of each rectrix were randomly scanned ten times each. Upon each use, the spectrometer was calibrated by scanning an Ocean Optics WS-1 white standard and a Colorline Ebony #142 dark standard. Additionally, the spectrometer was also calibrated between each feather using the Ocean Optics WS-1 white standard.

We quantified plumage colouration using traditional colourimetric values (e.g. Montgomerie 2006) and using tetrahedral models which correct for the avian visual system (e.g. Stoddard and Prum 2008; Fig. 2). Because nearly all studies on American redstarts use traditional colourimetric variables, we chose to include them in our study to allow comparison with the previous literature. To generate traditional color metrics of brightness, red chroma, and hue from Montogmerie (2006), we utilized the R-based program RCLR v.28. Reflectance spectra were imported into RCLR v.28 where we first performed smoothing using a LOESS function to eliminate noise and local peaks in the curves. Next, we calculated color variables of brightness, red chroma, and hue from the smoothed curves following Montgmerie (2006). Brightness (mean *R*_300–700_) was calculated as the mean amount of light reflected from 300–700 nm. Red chroma (*R*_605–700_/ *R*_300–700_) was measured as the proportion of light reflected in the orange-red segment of the spectrum relative to the overall light reflected. Hue (arctan ([(*R*_415–510_-*R*_320–415_)/*R*_320–700_]/ [(*R*_605–700_-R_400–500_)/*R*_320–700_])) was measured via segment classification to account for the proportion of light reflected in different regions of the spectrum (Saks et al. 2003). High hue values reflect dominance of shorter wavelengths (yellow-shifted), while low hue represent a dominance of longer wavelengths (red-shifted). Next, we averaged the colour variables from the ten reflectance spectra generated from each individual to obtain a single value of brightness, red chroma, and hue for each individual. These techniques follow those used in previous work by our group on American redstarts (Reudink et al. 2009a; Osmond et al. 2013; Tonra et al. 2014).

**Fig. 2.**
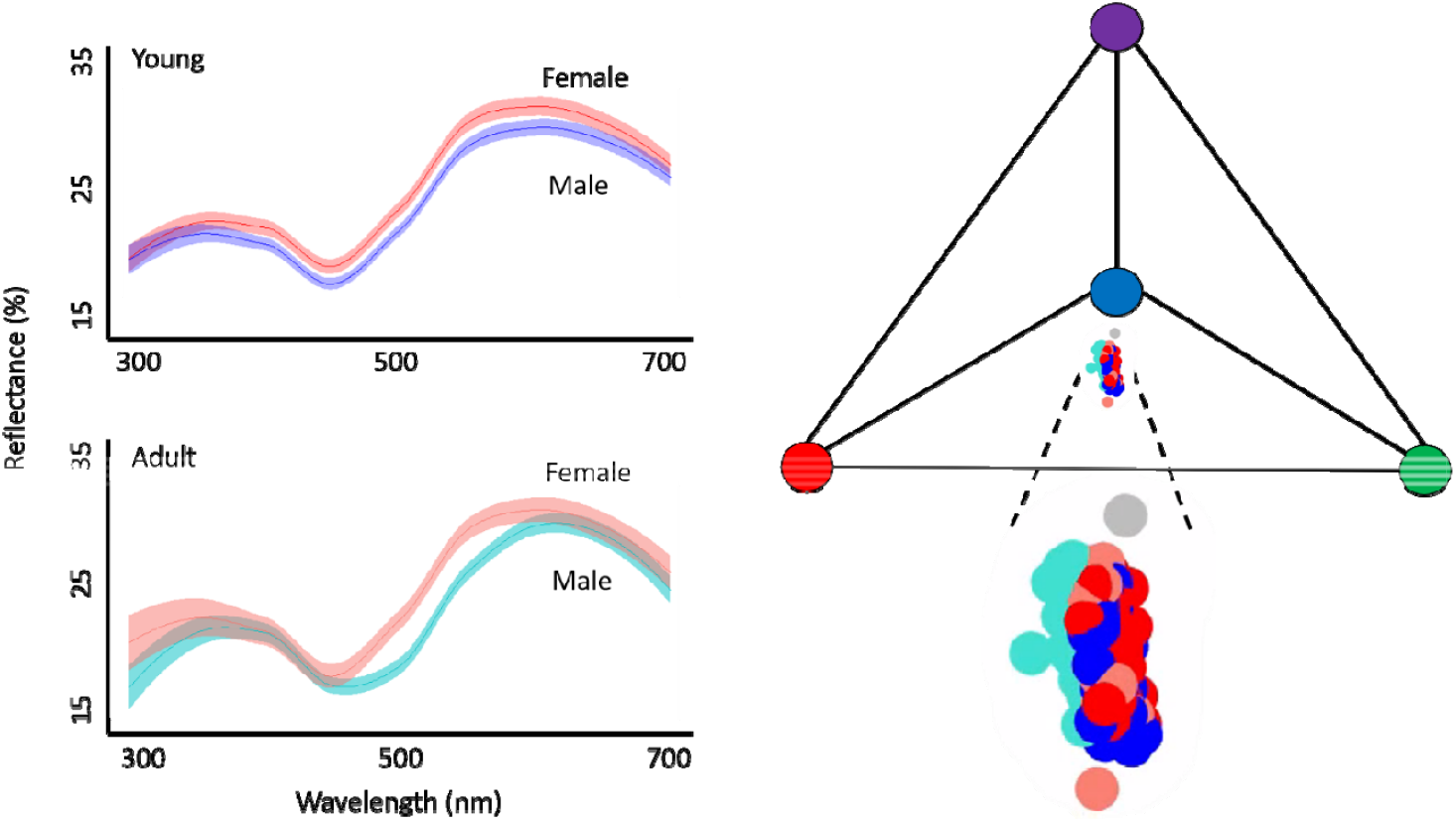
American redstart plumage colour varied by age and sex. Left panels: Reflectance plots showing the percentage (±SE) of light reflected off the feather surface. Right panel: Tetrahedral plot depicting age and sex plumage colour variation after correcting for the avian visual system. Grey dot represents the achromatic center of tetrahedral space.

We also quantified colour using tetrahedral models which correct for avian visual systems (Vorobyev et al. 1998), in the R package *pavo2* (Maia et al. 2019). Although the visual system of American redstarts is unknown, most passerines are ultraviolet sensitive (Ödeen and Håstad 2003), so our tetrahedral models use this visual system. We used a receptor-noise discrimination model to calculate the photon catch of each cone type used in avian colour vision. We used a value of 0.1 for colour (chromatic) and luminance (achromatic) discrimination and the photoreceptor ratios to 1:2:2:4 (ultraviolet, shortwave, mediumwave, longwave) in our models (Maier and Bowmaker 1993), which were selected based on a recent review of noise receptor estimates (Olsson et al. 2018). The models calculate hue θ, hue Φ, r.achieved, and luminance. Hue θ represents the angle of the color vector in the longitudinal plane while hue Φ represents the angle in the latitudinal plane and together hue θ and Φ describe the hue or the direction of colour vector in tetrahedral space (Stoddard and Prum 2008). r. achieved represents the chroma or saturation, or how different the colour is from the achromatic center of the tetrahedron (Stoddard and Prum 2008). Finally, luminance is analogous to brightness and represents the overall amount of light reflected from the feather sample and calculated independent of colour. Although this may seem redundant to include luminance, because all other previous redstart papers include traditional brightness, we thought it important to include both brightness and luminance in our study.

### Statistical Analyses

We tested how American redstart colour variation was explained by site, season, sex, age, and isotopic signatures at sampling using separate mixed models in the *lme4* package in R. In our models, we used traditional colourimetric variables (brightness, red chroma, and red hue) and colour modelled in tetrahedral space (hue θ, hue Φ, r.achieved, and luminance). In separate models, each colour variable was included as the response variable. Site, season, sex, age, δ^13^C and δ^15^N were main effects, and the interaction between age and season was included because redstarts exhibit delayed colour maturation (Hawkins et al. 2012). Year was included as a random effect. We then undertook model reduction for all mixed models based on the change in Akaike information criterion corrected for small sample sizes (AICc, Burnham and Anderson 2003) between the full model and each reduced model. We chose our final model based on the lowest AIC (Burnham and Anderson 2003). ΔAIC values within 4 were considered competitive and we selected the best fitting model using ANOVAs.

## Results

### Colour Variation

Our AICc model selection identified the reduced models for all response variables as the top models (Table 2 and Table S2). American redstart brightness was explained by site, season, and δ^15^N (Table 2). In both the spring and fall, American redstart feathers were brighter at the AIMS site (Fig. 3, site: F_1,134_=6.45, P=0.012; Season: F_1,134_=12.82, P<0.0001). Brightness was positively related to δ^15^N (Fig. 3. F_1,134_=11.78, P=0.001). Red chroma did not vary among any of our predictor variables (Table 2).

**Fig. 3.**
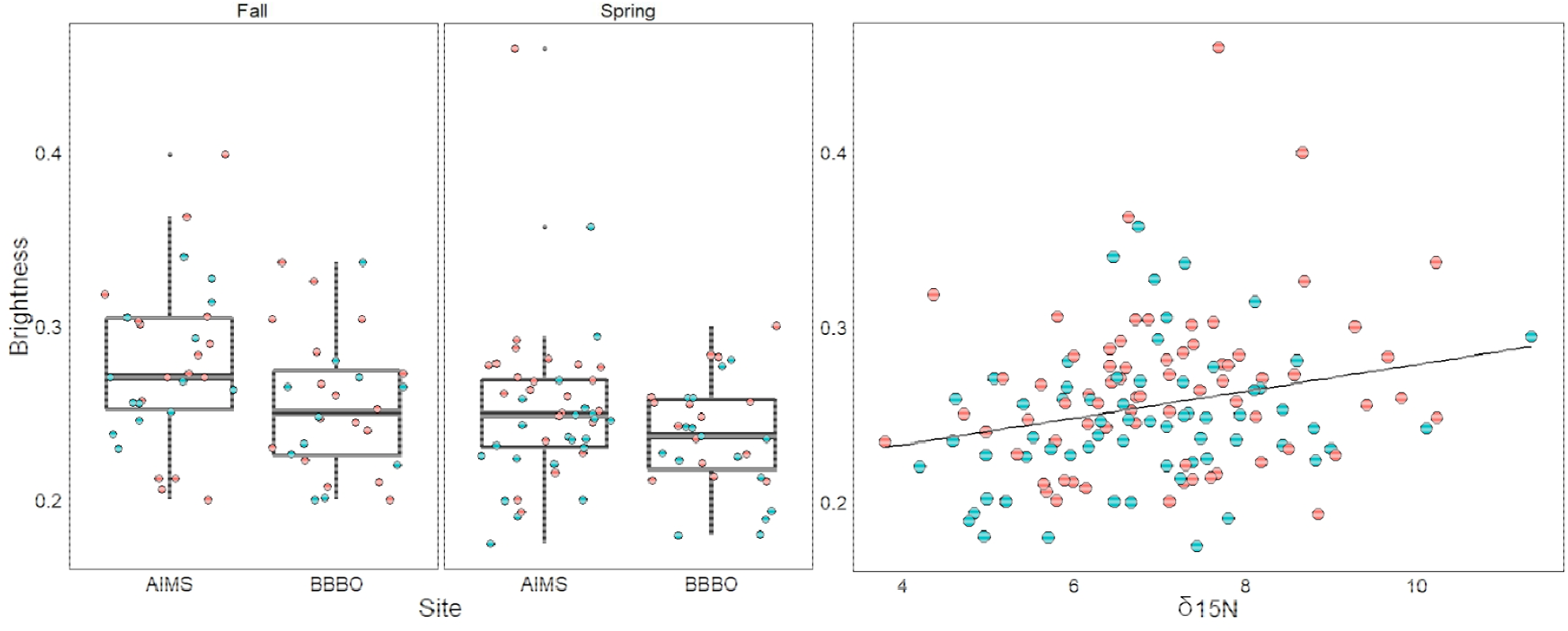
Left panels: American redstart feather traditional brightness (from Montgomerie 2006) in two sampling sites (AIMS and BBBO) in Fall and Spring. Right panel: Brightness and δ^15^N sampled from American redstart feathers. Females are indicated by red dots and males are indicated by blue dots.

**Table 2.** Model results demonstrating the effect of AICc selected fixed effects of site, season, sex, age, δ^15^N, and δ^13^C on American redstart colouration (brightness, red chroma, red hue, and tetrahedral colourspace hue θ, Φ, r.achieved, and luminance), and variances of the random effects of year. Significant effects are in bold.

| Variable | Fixed effect | Model *df* | Residual *df* | *F* | *P* | Random effect $\sigma^2$ |
| --- | --- | --- | --- | --- | --- | --- |
| Brightness | **Site** | **1** | **134.0** | **6.45** | **0.012** | 0 |
|  | **Season** | **1** | **134.0** | **12.82** | **<0.0001** |  |
|  | **$\delta^{15}\text{N}$** | **1** | **134.0** | **11.78** | **0.001** |  |
| Red Chroma | Site | 1 | 134.0 | 3.16 | 0.078 | 0 |
| Red Hue | **Season** | **1** | **134.0** | **26.09** | **<0.0001** | 0 |
|  | **Age** | **1** | **134.0** | **9.43** | **0.003** |  |
|  | **Sex** | **1** | **134.0** | **12.39** | **0.001** |  |
|  | **Season x Age** | **3** | **134.0** | **7.96** | **0.006** |  |
| Hue $\theta$ | **Site** | **1** | **133.9** | **5.93** | **0.02** | 0 |
|  | **Season x Age** | **3** | **133.9** | **30.11** | **<0.0001** |  |
|  | **Sex x Age** | **2** | **133.9** | **49.50** | **<0.0001** |  |
| Hue $\Phi$ | **Site** | **1** | **133.8** | **9.86** | **0.002** | 0.03 |
|  | **Season** | **1** | **133.6** | **129.14** | **<0.0001** |  |
|  | **Age** | **1** | **132.0** | **24.12** | **<0.0001** |  |
|  | **Sex** | **1** | **132.0** | **5.89** | **0.02** |  |
| r.achieved | **Season** | **1** | **134.0** | **33.69** | **<0.0001** | 0.002 |
|  | **Age** | **1** | **131.9** | **6.89** | **0.01** |  |
|  | **$\delta^{15}\text{N}$** | **1** | **131.9** | **6.86** | **0.01** |  |
| Luminance | **$\delta^{15}\text{N}$** | **1** | **134.0** | **5.59** | **0.02** | 0.002 |

Hue was explained by season, sex, age, and the interaction between season x age (Table 2 and Table S2). Hue was more red-shifted in males (Fig. 4, F_1,134_=12.39, P=0.001) and in adults (Fig. 4, F_1,134_=9.43, P=0.003) and following moult in the fall (Fig. 4, F_1,134_=26.09, P<0.0001). There was an interaction between season and age indicating that although hue in young birds did not vary between seasons, adult hue was more red-shifted in the fall (Fig. 4, F_1,134_=7.96, P=0.006).

**Fig. 4.**
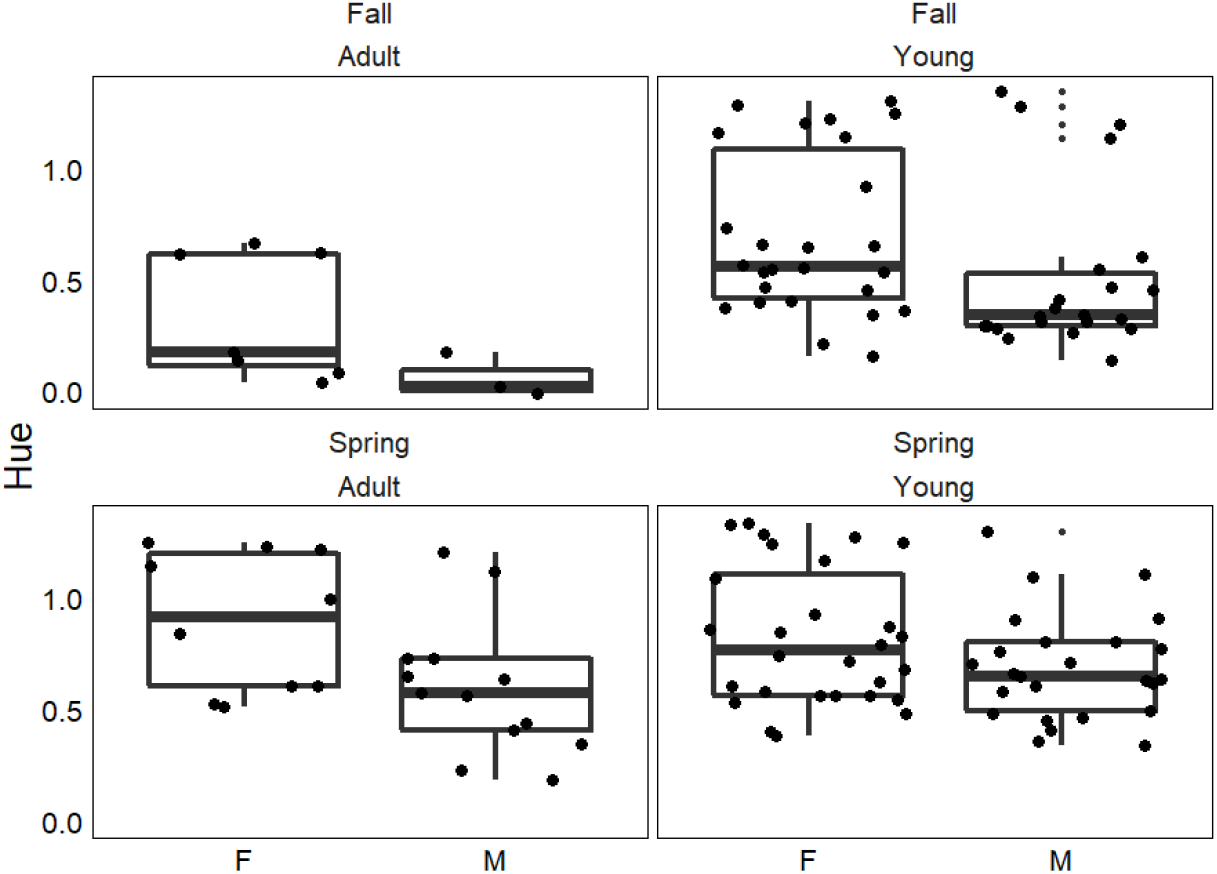
Female (F) and male (M) American redstart feather red hue (from Montgomerie 2006) in adult (left panels) and young (right panels) birds during fall (top) and spring (bottom) during fall (top) and spring (bottom).

Our analyses found redstart feather reflectance modelled in tetrahedral colourspace varied by several of our predictor variables (Table 2 and Table S2). Hue θ (Fig. 5) was predicted by site (F_1,133.9_=5.93, P=0.02) and the interactions between season x age (F_3,133.9_=30.11, P<0.0001) and sex x age (F_2,133.9_=49.50, P<0.0001). Hue Φ (Fig. 5) was predicted by site (F_1,133.8_=9.86, P=0.002), season (F_1,133.6_=129.14, P<0.0001), age (F_1,132.0_=24.12, P<0.0001), and sex (F_1,132.0_=5.89, P=0.02). Season (F_1,1340_=33.69, P<0.0001), age (F_1.131.9_=6.89, P=0.01), and δ^15^N (Fig. 6, F_1,131.9_=6.86, P=0.01) were significant predictors of r.achieved. Finally, luminance was predicted by δ^15^N (Fig. 6, F_1.134.0_=5.59, P=0.02).

**Fig. 5.**
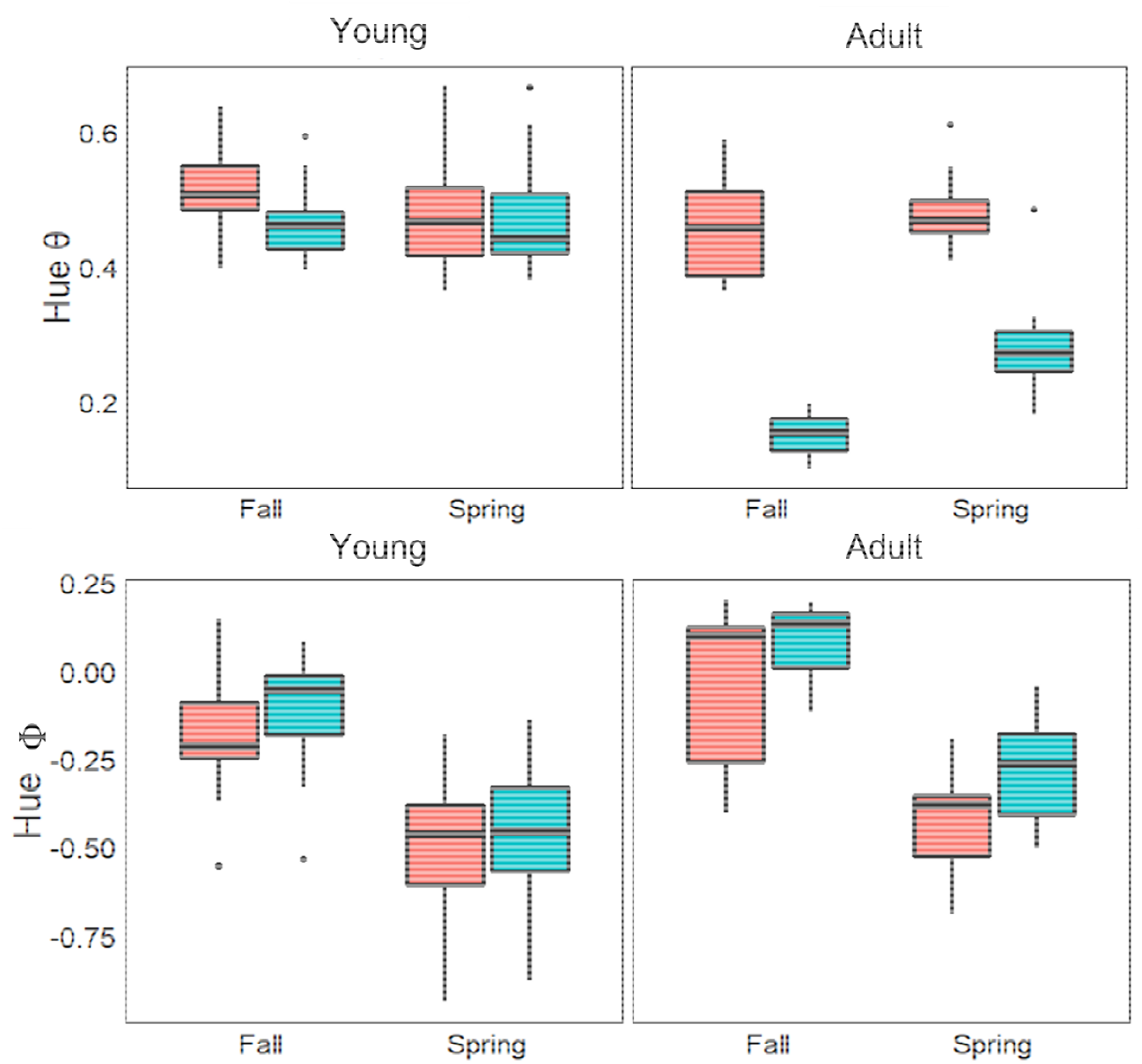
Female (red) and male (blue) American redstart tailfeather colour variation as modelled in tetrahedral colourspace. Hue θ represents the angle of the color vector in the longitudinal plane, while hue Φ represents the angle in the latitudinal plane. Together hue θ and hue Φ describe the hue or the direction of colour vector in tetrahedral space (Stoddard and Prum 2008). Top panel: hue θ varied by season, age, and sex. Bottom panel: Hue Φ similarly varied by season, age, and sex.

**Fig. 6.**
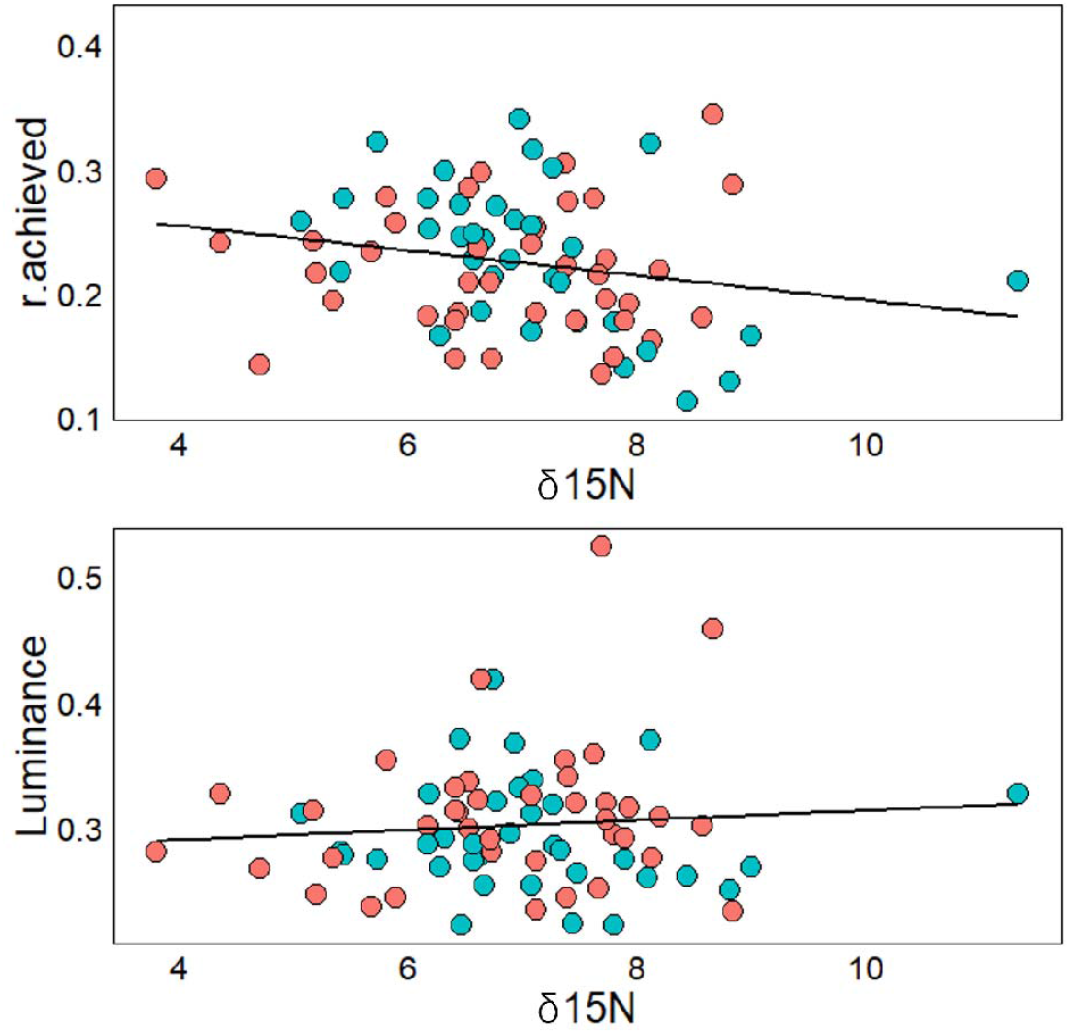
American redstart feather reflectance as modelled in tetrahedral colourspace (Stoddard and Prum 2008). Females are shown in red and males in blue. Top panel: r.achieved, representing the total length of the colour vector from the achromatic center of the tetrahedron, decreased with increasing δ^15^N. Bottom panel: Luminance, representing the total amount of light reflected from the feather sample increased with δ^15^N.

## Discussion

Carotenoid-based colouration serves important inter- and intra-sexual signaling functions; however, for some species like the American redstart, the relationship between fitness-related components and colour metrics can be perplexing and seemingly contradictory (Reudink et al. 2015). Part of the complexity likely stems from our incomplete understanding of the mechanisms underlying variation in American redstart colouration. Here, we demonstrate that geographic variation, season, and diet all underly intra-individual differences in American redstart colouration. However, the extent to which these factors influence colour expression is largely dependent on the colour metric under study, likely because different colour metrics reflect different aspects of feather quality (e.g., structural components vs. pigment deposition).

Geographic variation in plumage ornamentation has been widely documented across numerous species (Hill 2006; Badyaev et al. 2012) including the American redstart (Norris et al. 2007). Norris et al. (2007) found that individuals moulting at higher latitudes expressed plumage with higher red chroma compared to those at lower latitudes. In our study, we found differences in colouration between our two study sites (separated by 580km at similar latitudes) in three variables: brightness, hue θ, and hue Φ. All birds in our study were captured during migration and, as such, we lack information in the specific moult origins for each individual. Norris et al. (2007) suggested that geographic variation in American redstart colouration was likely driven by differences in dietary carotenoid availability or condition-related effects (e.g., parasite load) mediating carotenoid metabolism. Hue and chroma variables should be most closely linked to carotenoid content (Saks et al. 2003; Montgomerie 2006) and while brightness is negatively associated with feather carotenoid content, it is strongly mediated by feather structure (Shawkey and Hill 2005). Thus, while the site-specific differences we observed may be due to differences in carotenoid availability at the moult location, these differences may also reflect longitudinal geographic variation in condition, as one site was inland while the other was more coastal, but both were of similar latitude.

As expected, American redstarts expressed differences in brightness, hue θ, hue Φ, and r.achieved with higher brightness, and more red-shifted hue during the fall, following molt. However, variation in hue was dependent on age; while adults were more red-shifted in fall than spring, we detected no differences in first-year birds, potentially indicating that the orange colouration expressed by adult males may be more susceptible to the effects of UV degradation, feather-degrading bacteria, or feather wear between seasons (Örnborg et al. 2002; Delhey et al. 2006; Shawkey et al. 2007; Shawkey et al. 2009). Alternatively, if birds passing through migratory stopover sites in spring and fall differ in their moult location, differences between spring and fall plumage could simply reflect geographic variation in moult location.

Our brightness metrics (brightness and luminance) were associated with δ^15^N; birds with higher feather δ^15^N values had feathers with higher brightness. Because δ^15^N is directly linked to diet (Pearson et al. 2003; Gómez et al. 2018), these results suggest that American redstarts eating arthropods enriched with ^15^N (e.g., *Heteromurus nitidus, Drosophilia melanogastor*; Oelbermann and Scheu 2002) likely are not acquiring additional dietary carotenoids; however, these dietary sources may be advantageous during feather growth, allowing the birds to create feathers with a high-quality microstructure. Whether or not feather brightness and luminance *per se* is a signal used by American redstarts is unclear; however, Reudink et al. (2009b) demonstrated that males that occupied higher-quality winter-territories expressed feathers with higher brightness and Reudink et al. (2009a) found that polygynous males had higher feather brightness than monogamous males. Because polygyny requires the ability to hold multiple territories or a single large territory, Reudink et al. (2009a) suggested that feather brightness may act as a social signal during both the breeding and wintering season, directly reflecting a male’s competitive ability. Thus, diet during moult may potentially impact intrasexual signaling.

In a study of Bullock’s orioles, Sparrow et al. (2017) found that feather δ^15^N values were associated with feather colouration and feather carotenoid content. The authors found that while δ^15^N values were positively associated with overall carotenoid concentration, birds with more orange-shifted hue had feathers with lower δ^15^N values. The authors suggested that low δ^15^N values could indicate a diet with a higher fruit:insect ratio and birds with more orange-shifted hue, though they had lower overall carotenoid concentrations, may have had a higher abundance of specific orange or red carotenoids. It is important to note that American redstarts and Bullock’s oriole diets differ considerably; orioles have a more varied diet of both fruits and insects (Flood et al. 2016), while redstarts are primarily insectivores (Robinson and Holmes 1982). Thus, the way in which δ^15^N is associated with the expression of carotenoid-based colouration is likely highly dependent on the dietary ecology of the species. It is also important to consider that our observed differences may be due to accumulation of dirt in feathers or due to degradation caused by UV exposure (Örnborg et al. 2002). The decrease in brightness and hue between the fall and spring is consistent with previous studies (Prum et al. 1999). Interestingly, while our tetrahedral metric for chroma (r.achieved) did vary in our study, our traditional colourimetric variable (red chroma) did not significantly vary, as was previously found in carotenoid-red colouration as a result of abrasion or soiling (McGraw and Hill 2004). However, without estimates of dirt on feathers, microscopy of feather nanostructure, or repeated measures, we can only speculate on this effect and encourage future work to consider adventitious colouration.

## Notes

### Competing Interest Statement

The authors have declared no competing interest.

### Summary of Updates

Changes to the manuscript appear in red.

